# PCNA Inhibition Enhances the Antitumor Activity of KRAS-Targeted Therapies in Pancreatic Cancer

**DOI:** 10.64898/2025.12.06.692635

**Authors:** Sahar F Bannoura, Husain Yar Khan, Amro Aboukameel, Yin Wan, Bin Bao, Md Hafiz Uddin, Hugo Jimenez, Adeeb Aboukameel, Rayyan Siddiqui, Filza Khan, Pouya Haratipour, Long Gu, Rafic Beydoun, Muhammad Wasif Saif, Miguel Tobon, Philip A. Philip, Bassel El-Rayes, Herbert Chen, Robert J Hickey, Linda Malkas, Yang Shi, Mohammed Najeeb Al Hallak, Ramzi M Mohammad, Boris C. Pasche, Asfar S. Azmi

**Affiliations:** Department of Oncology, Wayne State University School of Medicine, Barbara Ann Karmanos Cancer Institute, Detroit, MI, USA; Beckman Research Institute of City of Hope, Duarte, California, USA; Department of Pathology, Wayne State University School of Medicine, Detroit, MI, USA; Henry Ford Cancer Institute - Henry Ford Health, Detroit, MI, USA; Department of Hematology and Oncology, University of Alabama at Birmingham, O’Neill Comprehensive Cancer Center, Birmingham, AL, USA; Department of Surgery, University of Alabama at Birmingham, Birmingham, AL, USA

## Abstract

Pancreatic ductal adenocarcinoma (PDAC) is an aggressive disease with a dismal prognosis. More than 90% of PDAC tumors harbor KRAS mutations, and several KRAS inhibitors, such as off-state, on-state, mutation-specific, and pan-RAS inhibitors, are being tested in preclinical and clinical settings. However, the response to these inhibitors as single agents is less than optimal, indicating the need to identify novel combination therapies to improve treatment outcomes. Proliferating cell nuclear antigen (PCNA) is a ring-shaped clamp protein that regulates DNA replication, repair, and resolution of transcription-replication conflict, which are critical processes for pancreatic cancer survival. AOH1996 is a first-in-class, selective PCNA inhibitor in Phase I trials. Here, we found that AOH1996 treatment is efficacious in various PDAC models in vitro. PCNA and KRAS are predicted to be synthetic lethal partners, and RNA sequencing of AOH1996-treated PDAC cells reveals enrichment of MAPK and PI3K signaling pathways. Combination of AOH1996 with KRAS inhibitors demonstrates strong synergy across KRAS G12C and G12D mutant models. Treatment with a combination of AOH1996 and KRAS inhibitors induces cell cycle arrest and apoptosis in PDAC cells. Robust antitumor activity of AOH1996 in combination with RMC-6236 was observed in PDAC tumoroids. *In vivo*, the combination of AOH1996 with sotorasib or MRTX1133 reduced tumor growth rates compared to single-agent therapy, with no impact on mouse body weight. Residual tumor analysis showed sustained pERK and Myc inhibition in the combination arm. In conclusion, combination of AOH1996 with KRAS inhibitors is a promising therapeutic strategy for KRAS-driven PDAC, warranting further clinical investigation.

## Introduction

Pancreatic ductal adenocarcinoma (PDAC) is an aggressive disease with a dismal prognosis. It is the 9^th^ most common cancer and the 3^rd^ leading cause of mortality in cancer patients in the United States^1^. With an increasing incidence rate, PDAC poses a significant clinical challenge with advanced disease at diagnosis, poor response to therapy, and common recurrence^2,3^.

KRAS mutations are present in many different solid and hematological tumors. In PDAC, around 90% of patients have oncogenic KRAS mutations^4^. Several KRAS inhibitors have emerged recently, including mutation-specific on- and off-state inhibitors as well as pan-RAS inhibitors. These KRAS inhibitors have been shown to block downstream KRAS signaling, thus restricting tumor growth^5-7^. Nonetheless, acquired resistance to these inhibitors is a significant clinical problem^8,9^; therefore, efficacious therapeutic combinations are crucial for better responses and avoiding resistance^10^.

Proliferating cell nuclear antigen (PCNA) is a ring-shaped protein that acts as a sliding clamp during DNA replication and has essential functions in regulating DNA repair^11,12^. In cancer, PCNA has various tumor-promoting roles. A high level of PCNA expression is a hallmark of rapid proliferation, aggressive tumor behavior, and poor prognosis. Additionally, PCNA’s role in DNA repair helps cancer cells tolerate DNA damage, thus promoting the survival of cancer cells with high DNA damage. Central to the multifunctional roles of PCNA are its various posttranslational modifications, including ubiquitination, SUMOylation, phosphorylation and acetylation. These PTMs help fine-tune the function of PCNA to respond to different DNA damage or DNA replication stress stimuli by promoting different repair pathways. A PTM on the interdomain connector loop of PCNA has been identified as cancer-associated and as a potential cancer-specific therapeutic target^13,14^. Exploiting this cancer-specific vulnerability is a novel small molecular inhibitor of PCNA, known as AOH1996, that binds to its interdomain connector loop and prevents its interaction with p21 and TLS polymerases^14,15^. AOH1996 is currently in phase I/II clinical trials for solid tumors.

In the present study, we demonstrate the efficacy of AOH1996 in PDAC and examine the underlying molecular mechanisms. We show that AOH1996 synergizes with various KRAS inhibitors using cell culture, patient-derived xenografts, and *in vivo* assays.

## Materials and Methods

### Cell culture

The pancreatic cancer cell lines MIA PaCa-2 (RRID:CVCL_0428), HPAC (RRID: CVCL_3517), PANC-1 (RRID:CVCL_0480) cells were maintained in DMEM media (Gibco), HPAF-II(RRID:CVCL_0313) in EMEM media (Gibco), AsPC-1 (RRID: CVCL_0152) in RPMI-1640(Gibco). Media were supplemented with 10% FBS and 1% penicillin/streptomycin (Gibco). Cells were maintained at 37°C in a humidified 5% CO_2_ atmosphere. Cell lines have been authenticated at a core facility at Wayne State University (WSU), using short tandem repeat (STR) profiling on the PowerPlex 16 System (Promega). The cell lines were routinely tested for Mycoplasma contamination using PCR.

### Compounds

Sotorasib (AMG 510), adagrasib (MRTX849), MRTX1133, and RMC-6236 were purchased from Selleck Chemicals LLC (Houston, TX) or BOC Sciences (Shirley, NY). Oxaliplatin was obtained from Karmanos Cancer Institute (KCI) pharmacy. All drugs were prepared as 50mM stock solutions in cell culture grade DMSO, 10mM working stock solution aliquots, and freshly diluted in the appropriate media immediately before use for each experiment.

### Cell Viability Assay and Synergy Analysis

Depending on the cell line, 2500 to 4000 cells were seeded in 96-well microtiter culture plates at equal densities and allowed to grow overnight at 37°C. Media was then replaced with treatment media containing vehicle or test compounds at a range of concentrations alone or in combination for 72 hours for single-agent IC_50_ determination or 120 hours for combination synergy determination using a 6 x 6 matrix. Cell viability was assessed using an MTT assay. The reagent 3-(4,5-dimethylthiazol-2-yl)-2,5-diphenyltetrazolium bromide (MTT) was added to the wells at a final concentration of (0.5 mg/mL) and incubated for 3 hours at 37°C. After the incubation period, the reaction was terminated by aspirating the supernatant and adding DMSO to solubilize the formazan crystals. After 15 minutes of gentle shaking, the absorbance was measured at 570nm on a SynergyHT plate reader (BioTek, Winooski, WI, USA). Relative viability was calculated using absorbance data where all individual readings were normalized to average vehicle control and then analyzed and plotted using GraphPad Prism 10 (RRID:SCR_002798). Combination Synergy calculations were performed using SynergyFinder+ online software tool (RRID:SCR_019318).

### Patient-derived PDAC tumoroid model and 3D CTG assay

Human PDAC tumoroid cultures were developed from surgically resected de-identified primary tumor tissue specimens under an active IRB protocol (#034916MP2X). The tumoroids, consisting of both cancer-associated fibroblasts and tumor cells, were cultured in RPMI media and maintained in 3D conditions using ultra-low attachment plates. These PDAC tumoroids were treated with 1 μM AOH1996 or 100 nM RMC6236 or their combination, twice a week. After one week of drug treatment, tumoroids were imaged using an inverted phase-contrast microscope fitted with a camera (Olympus). The effect of drug treatments on the viability of tumoroids was quantified using 3D CellTiter-Glo (CTG) luminescent cell viability assay (Promega cat#G7570) according to the manufacturer’s protocol.

### Western Blotting

Total protein extracts were prepared by lysing the cells in ice-cold RIPA buffer (Santa Cruz Biotechnology; SCBT cat#sc-24948) supplemented with protease and phosphatase inhibitors. Protein concentration was measured using Pierce BCA protein assay (Thermo Scientific cat#23225). Samples were prepared in 6x Laemmli’s buffer (Thermo Scientific cat#J61337.AD) and boiled for 5 minutes. The protein samples were separated using 4-20% SDS-PAGE Mini-PROTEAN TGX Precast Gels (Bio-Rad cat#4561093) at 100V. Gels were subsequently transferred to an Immun-Blot PVDF membrane (Bio-Rad #162-0177) using the Mini Trans-Blot wet transfer system at 100V for 1 hour. After transfer, membranes were blocked using EveryBlot Blocking Buffer (Bio-Rad cat#12010020) for 10 minutes at room temperature and then incubated in a specific primary antibody overnight at 4°C; specific conjugated secondary antibodies were added for 1 hour at room temperature (RT). The signal was detected by chemiluminescence using SuperSignal West Pico PLUS substrate (Thermo Scientific cat#34580) on a Bio-rad ChemiDoc MP Imaging System (RRID:SCR_019037). ImageJ Software (RRID:SCR_003070) was used for densitometric quantification. The following antibodies were used: GAPDH, source: SCBT: cat#sc-365062, 1:2500 dilution. Ran, source: Cell Signaling Technology (CST) Cat# 4462 1:1000 dilution. Phospho-AKT (pAKT, Ser473), source: SCT cat#4060, 1:1000 dilution. AKT source: SCT cat#9272, 1:1000 dilution. Phosphor-p44/p42 MAPK (phospho-ERK1/2, pERK) (Thr202/204), source: SCT cat#9101, 1:1000 dilution. P44/42 MAPK (ERK1/2), source: SCT cat#4695, 1:1000 dilution. Anti-mouse IgG, HRP-linked Antibody, source: SCT cat#7076, 1:3000 dilution. Anti-rabbit IgG, HRP-linked Antibody, source: SCT cat#7074, 1:3000 dilution.

### RNA extraction and Real-Time quantitative PCR

Total RNA was purified from cells using the RNeasy Plus Mini Kit (Qiagen, cat#74134), according to the manufacturer’s protocol. RNA was reverse transcribed using the High-Capacity cDNA Reverse Transcription Kit (Applied Biosystems cat#4368814), and mRNA expression was analyzed by real-time qPCR using SYBR Green Master Mix (Applied Biosystems, cat# A25743). The qPCR was initiated by 10 min at 95°C, followed by 40 thermal cycles of 15 s at 95°C and 1 min at 60 °C in a StepOnePlus real-time PCR system (Applied Biosystems). Table S1 lists primers used.

### RNA sequencing and bioinformatics analysis

Poly-A RNA sequencing was performed at Novogene. Messenger RNA was purified by poly-T oligo attached magnetic beads from total RNA. After fragmentation, the first strand cDNA was synthesized using random hexamer primers, followed by the second strand cDNA synthesis using either dUTP for the directional library or dTTP for the non-directional library. For the non-directional library, it was ready after end repair, A-tailing, adapter ligation, size selection, amplification, and purification. For the directional library, it was ready after end repair, A-tailing, adapter ligation, size selection, USER enzyme digestion, amplification, and purification. The library was checked with Qubit and real-time PCR for quantification and bioanalyzer for size distribution detection. Quantified libraries were pooled and sequenced on Illumina platforms, according to effective library concentration and data amount. The raw Fastq files were quantified using kallisto (v0.50.0) with the transcriptome indices built on the cDNA FASTA files from human (Homo sapiens) genome assembly GRCh38. Differential gene expression was analyzed using DESeq2 (v1.50.2), and the differentially expressed genes (DEGs) were identified using the thresholds of |log2(foldchange)| > log2(1.5) and a false discovery rate (FDR) < 0.05. Functional enrichment analysis was performed using clusterProfiler (v4.18.2) with the Kyoto Encyclopedia of Genes and Genomes (KEGG) pathway database.

### Flow cytometric apoptosis analysis

Cells were sorted using Annexin V FITC and Propidium Iodide (PI) according to the manufacturer’s protocol. After treatment, cells were dissociated with Trypsin-EDTA and resuspended in media and allowed to recover at 37°C for 20 minutes to regain cell membrane integrity. The cells were then stained with Annexin V and PI. Stained cells were sorted using the Cytek Northern Light flow cytometer at the WSU Microscopy, Imaging and Cytometry Resources (MICR) Core.

### Cell cycle assay

Cell cycle phase was determined using DNA content analysis by flow cytometry. Cells were harvested and fixed in ice-cold 70% ethanol, and left to fix overnight at 4°C. The following day, the cells were washed twice in PBS, then permeabilized with Triton X-100 and stained using PI with DNase-free RNase. DNA content was detected using Cytek Northern Light flow cytometer at the WSU MICR Core. Data was analyzed using ModFit LT (RRID:SCR_016106).

### Clonogenic assay

Cells were counted and reseeded (500 cells/well) in a 6-well plate and allowed to adhere overnight. The following day, cells were treated with AOH1996, KRAS inhibitors, their combination, or vehicle control. Fourteen days later, the wells containing the colonies were washed with PBS, fixed with methanol, and stained with crystal violet for 30 minutes. The plates were then carefully washed in tap water, dried, and photographed. Colonies were manually counted and depicted as bar graphs.

### Spheroid formation assay

A single cell suspension (1000 cells/100 μL) was plated in 96-well ultra-low attachment plates (Corning) in 3D tumorsphere media XF (PromoCell). Media were replenished every 3 days and growth was monitored microscopically. After spheroids were formed, they were photographed, and viability quantified using CellTiter-Glo Luminescent Cell Viability Assay (Promega cat#G7570).

### *In vivo* mouse xenograft models

*In vivo* studies were conducted under WSU’s Institutional Animal Care and Use Committee (IACUC)-approved protocol #25-02-7587 in accordance with the approved guidelines.

AsPC-1 cell-derived tumor xenograft model:

AsPC-1 cells were washed in PBS and then resuspended in cold PBS at a concentration of 1 x 10^6^ cells per 100 μL. Donor ICR-SCID mice, aged 4–5 weeks, were inoculated unilaterally in the left flank using a BD 26Gx 5/8 1 mL Sub-Q syringe. Once the tumors reached approximately 5%–10% of body weight, the donor mice were euthanized, and the tumors were harvested. Tumor fragments were subsequently implanted into the left flank of recipient mice unilaterally. Seven days after transplantation, mice were randomized into four groups of six mice each. Compounds were orally administered as follows: MRTX1133 at 15 mg/kg daily Monday through Friday for 4 weeks; AOH1996 at 100 mg/kg twice daily Monday through Friday for 4 weeks; or a combination of the two treatment regimens. Tumor volumes were determined using the formula0.5 × *L* × *W*^2^ in which *L* refers to length and *W* refers to width of each tumor. Upon completion of drug dosing, tumor tissues from the control or treatment groups were harvested and analyzed used for IHC and Western blot.

MIA PaCa-2 cell-derived tumor xenograft model:

Tumors were established in the same manner described for AsPC-1 xenografts. Mice were randomized into 4 groups of 8 mice each and received oral gavage treatment. Treatment regimens were as follows: Vehicle treatment; or AOH1996 at 100 mg/kg twice daily for 5 days a week for 4 weeks, or sotorasib at 25 mg/kg 5 days a week for 4 weeks, or a combination of the two regimens. To perform survival analysis, mice were euthanized at a humane timepoint (up to 20% body weight), and survival was depicted using Kaplan-Meier Analysis.

### Immunohistochemistry (IHC)

IHC was performed on formalin fixed paraffin embedded (FFPE) at the Biobank and Correlative Sciences (BCS)_Core facility at Karmanos Cancer Institute, WSU, using a standard procedure. Mouse xenograft tissues were processed and stained with Eosin and Hematoxylin stains (Leica Biosystems) or immunohistochemistry (IHC) with one of the following antibodies, Ki67 (Source: Agilent Dako, Cat#M724029-2, dilution of 1:75), Phosphor-p44/p42 MAPK (phospho-ERK1/2;Thr202/204; source: SCT cat# 9106, dilution of 1:50), pAKT (source: SCT cat# 3787, dilution of 1:50). Heat induced epitope retrieval (HIER) was performed with a pH 6 buffer for 20minutes at 95°C, followed by endogenous peroxidase block using 3% hydrogen peroxide for 5 minutes at RT. Novolink Polymer Detection Systems kit (Leica) was used to develop the staining. H scoring was done with HALO v4.1 (RRID:SCR_018350).

### Statistical analysis

All experiments were performed in three replicates unless otherwise noted. P values (*, P < 0.05; **, P < 0.01, *** P < 0.001, *** P <0.0001) were determined by two-sided unpaired t-tests, unless otherwise noted. Multiple comparisons were corrected using a False Discovery Rate (FDR) correction of 5%. The distributions of survival outcomes were graphically summarized using Kaplan-Meier plots and comparisons between groups were conducted using a log-rank test.

## Results

### PCNA as a target for Pancreatic Ductal Adenocarcinoma

To determine the extent to which PCNA is involved in PDAC tumor maintenance and progression, we used data from GTEx and TCGA to compare expression levels in patient versus normal tissues, as well as correlation to prognosis. PDAC tumors expressed significantly higher levels of PCNA compared to normal pancreas **(Fig. 1A)**. An analysis of patient survival showed that those with higher PCNA expression had worse overall survival, progression-free survival, and disease-free survival **(Fig. 1B)**. We tested the efficacy of the novel PCNA inhibitor AOH1996 as a single agent in various PDAC models. PDAC cells showed a dose-response sensitivity to PCNA inhibition in the presence of AOH1996 **(Fig. 1C)**. PDAC cells also showed a dose-dependent sensitivity to AOH1996 in three-dimensional (3D) spheroid cultures **(Fig. 1D)**. Additionally, PCNA inhibition decreased the potential of PDAC cells for long-term proliferation and self-renewal as shown by a colony formation assay in a dose-response manner **(Fig. 1E)**. Notably, PDAC cells responded to AOH1996 treatment regardless of molecular classification. Cells with various mutations, including KRAS^G12D^ mutant cells (PANC-1, HPAC, AsPC-1), or KRAS^G12C^ mutated cells (MIA PaCa-2), efficiently respond to AOH1996 treatment. Additionally, cells considered to represent classical molecular subtype (AsPC-1, HPAC) as well as those classified as basal-like (MIA PaCa-2, PANC-1) were uniformly sensitive to PCNA inhibition.

**Figure 1.**
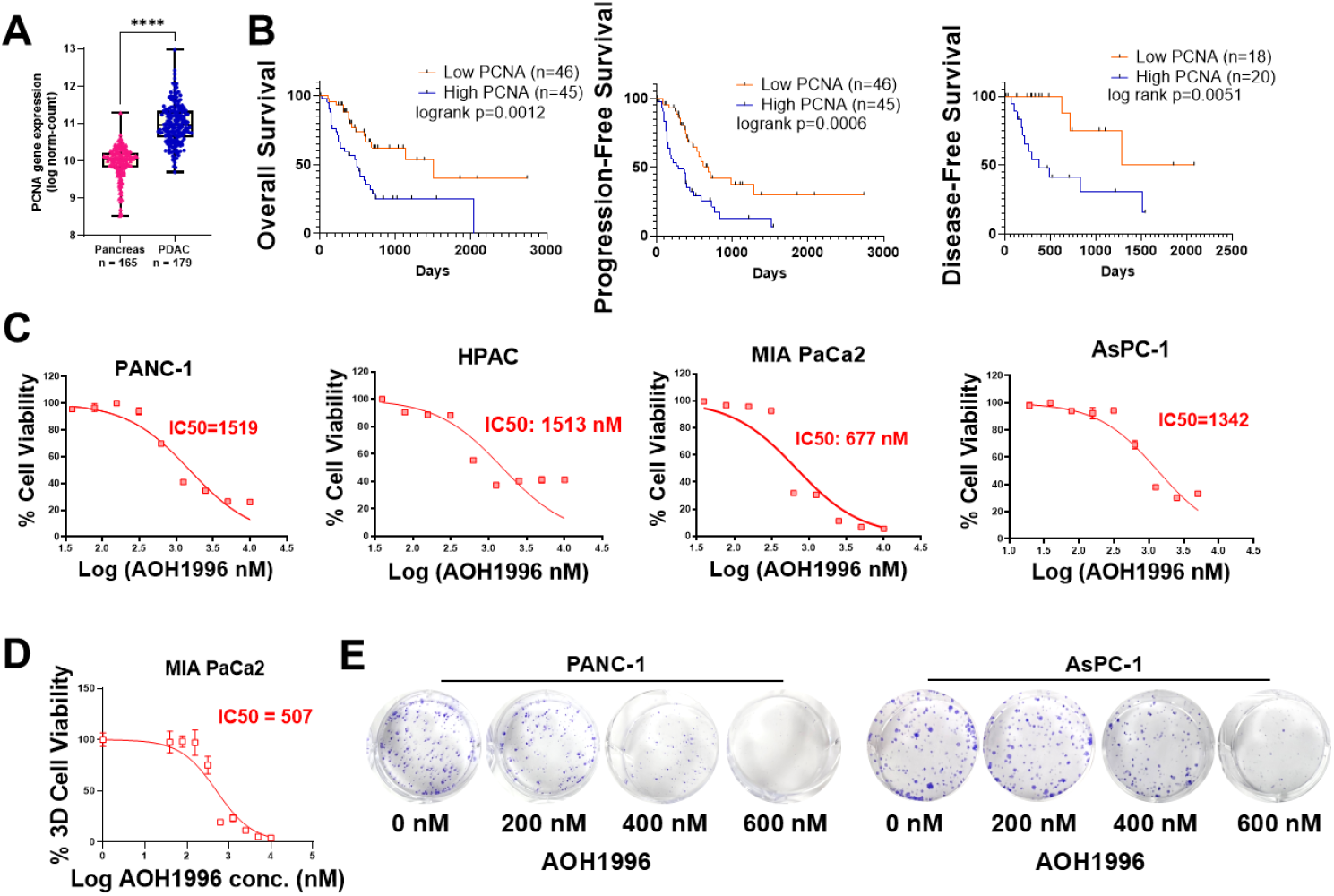
Novel PCNA inhibitor AOH1996 inhibits PDAC models *in vitro*. **(A)** PCNA expression levels in normal pancreas (GTEx) and PDAC tissues (TCGA). **(B)** Kaplan-Meier TCGA patient survival analysis based on PCNA levels showing overall survival, progression-free survival, and disease-free survival in high and low PCNA quartiles. **(C)** Dose response plots and IC50 estimation for AOH1996 in a panel of PDAC cell lines using MTT cell viability assay. **(D)** Dose response plot and IC50 estimation of MIA PaCa-2 derived spheroids using CellTiter Glo assay. **(E)** Colony formation assay of PDAC cells in the presence of increasing concentrations of AOH1996.

### PCNA inhibition causes broad perturbations in MAPK signaling and cancer-supporting pathways

To investigate the molecular features of PCNA inhibition in PDAC, we performed RNA sequencing in two different PDAC cell lines treated with AOH1996 or a vehicle control. The cells were treated for 48 hours, then RNA was extracted for polyA RNA sequencing. This analysis revealed significant transcriptional alterations, with many differentially expressed genes (DEGs) in both cell lines: 1,084 total DEGs in AsPC-1 cells **(Fig. 2A)** and 2,579 total DEGs in PANC1 cells **(Fig. 2B)**. Principal component analysis (PCA) showed that most of the variance in the dataset was represented by the identity of the cell line, followed by treatment with AOH1996 versus control **(Fig. S1A)**. This indicates that the treatment has a significant impact on the transcriptome of these cells. Heatmaps of the DEGs showed treatment samples clustering closely together and starkly separated from control samples, indicating high reproducibility of the results across replicates **(Fig. S1B**,**C)**. We validated sequencing results using RT-qPCR, which replicated RNA-seq data **(Fig. S1D)**. To find the pathways that are commonly regulated by PCNA inhibition, we examined the overlap of DEGs among the two cell lines, which revealed that around 10% of all DEGs were commonly impacted **(Fig. 2C)**, including 146 genes that were upregulated and 148 that were downregulated in both cell lines **(Fig. 2D)**. To understand pathway-level implications of gene regulation we performed Kyoto Encyclopedia of Genes and Genomes (KEGG) pathway enrichment analysis, thus revealing several perturbed pathways related to cancer signaling pathways and apoptosis **(Fig. 2E)**. Interestingly, several pathways were enriched in both datasets including Mitogen-Activated Protein Kinase (MAPK) signaling pathway, cellular senescence, PI3K-AKT and Hippo signaling pathways (**Fig. 2F**). MAPK pathway is an essential cellular signaling cascade that is of critical importance for PDAC cell proliferation^16^. PDAC tumors are addicted to the oncogene KRAS-activated ERK pathway for their survival and growth. Additionally, the PI3K-AKT pathways is one of several downstream pathways that KRAS may be able to activate and work in concert to drive cancer proliferation^17^. Therefore, we examined whether KRAS is a potential therapeutic co-target with PCNA. Using the Slorth database, which ranks potential synthetic lethal interactors by identifying and exploiting conserved patterns in protein interaction network topology both within and across species^18^, we found that KRAS is predicted to be one of the main synthetic lethal targets with PCNA **(Fig. S1E)**. Therefore, we decided to test the combination of AOH1996 with novel KRAS inhibitors.

**Figure 2.**
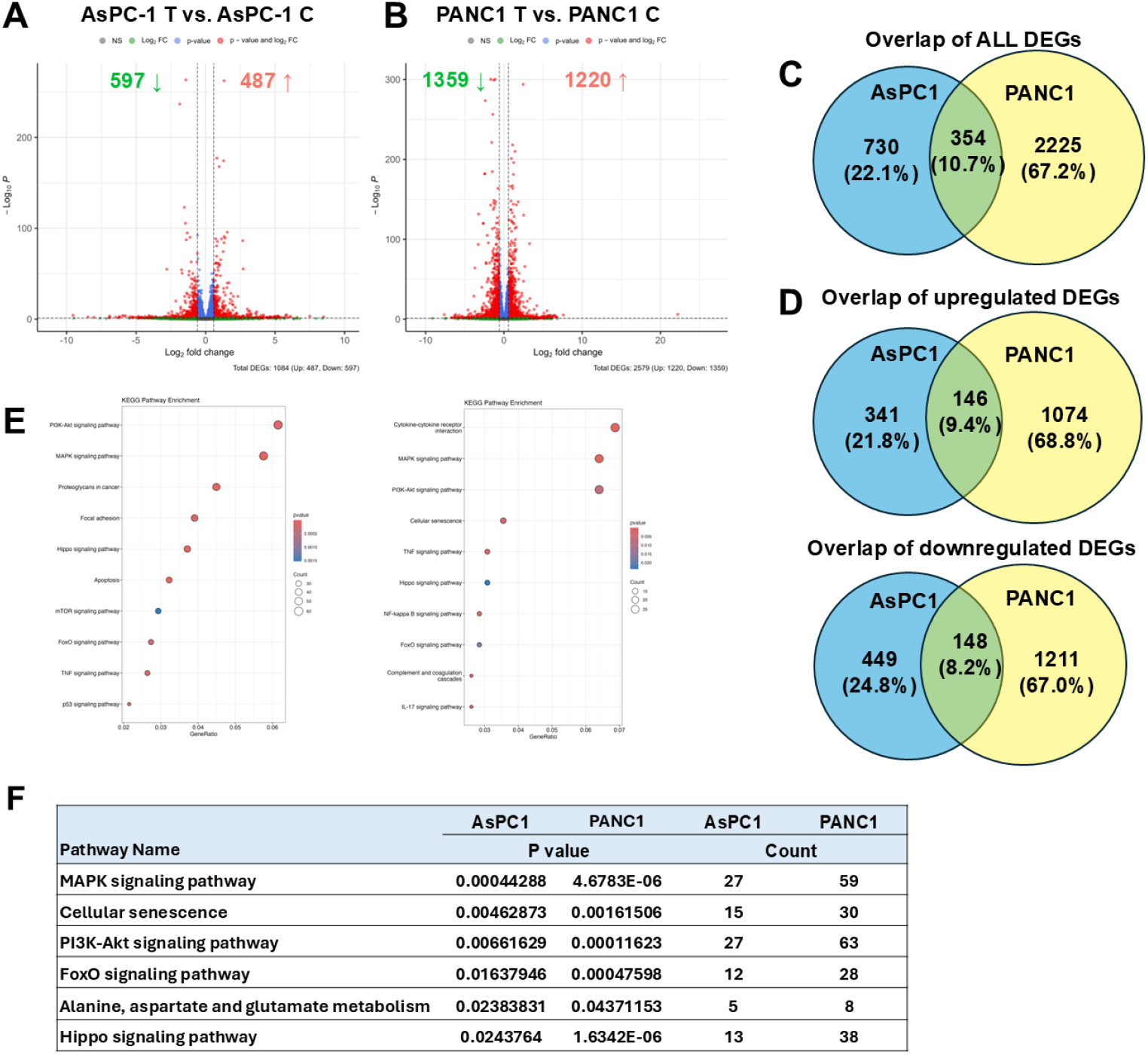
AOH1996 treatment induces widespread transcriptional alterations in PDAC cellular models impacting MAPK signaling. RNA sequencing reveals the differentially expressed genes (DEG) in response to AOH1996 treatment in two different PDAC cell lines. **(A)** Volcano plot depicting DEGs in AsPC-1 cell line. **(B)** The same as (A) in PANC1 cell line. **(C)** Venn diagram showing the overlap in DEGs between the two cell lines. **(D)** The same as (C) showing upregulated or downregulated DEG overlap. **(E)** KEGG pathway enrichment analysis of significant DEGs shown in (A) and (B). **(F)** Commonly enriched KEGG pathways are presented in the table.

### AOH1996 synergizes with KRAS^G12D^ and KRAS^G12C^ inhibitors *in vitro* and drives tumor regressions *In Vivo*

To find the relationship between AOH1996 and KRAS small molecule inhibitors in PDAC cells, we performed *in vitro* testing of a range of concentrations combining AOH1996 with MRTX1133, a KRAS^G12D^ inhibitor, and combinations of AOH1996 with AMG-510, or MRTX849, KRAS^G12C^ inhibitors, and calculated synergy scores using SynergyFinder^19^. Significant synergy was found at various dose combinations in three KRAS^G12D^ cell lines with MRTX1133 **(Fig. 3A)**. Colony formation assay also showed significant long-term inhibition of cancer growth *in vitro* in the combination **(Fig. 3B)**. The combination treatment also showed pERK and pAKT pathway inhibition at 24 hours **(Fig. 3C)**. Cell cycle analysis using DNA content staining followed by flow cytometry revealed a G2/M phase arrest in AOH1996 treated cells, and a G1 phase arrest in the MRTX1133 treated cells **(Fig. 3D)**. Combination-treated cells showed a significant increase in apoptotic cells, and a significant decrease in the cells undergoing replication in S phase with cells being arrested in G1 and G2/M phases **(Fig. 3D)**.

**Figure 3.**
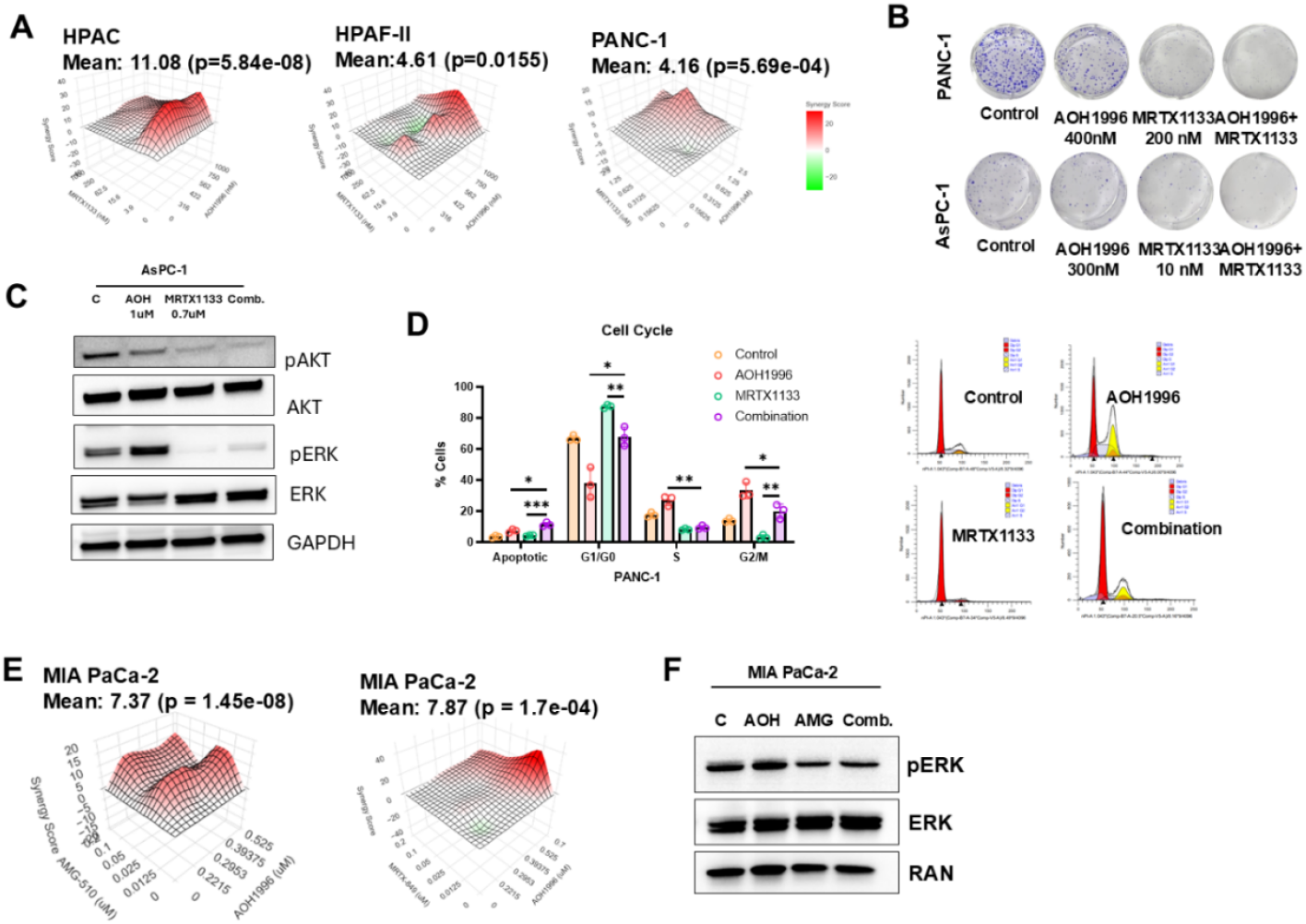
Combination of mutant KRAS inhibitors with PCNA inhibitor AOH1996 is synergistic in PDAC cell lines. **(A)** Synergy plots between AOH1996 and MRTX1133 in three different KRAS-driven PDAC cellular models. **(B)** Picture of a colony formation assay in PDAC cells treated with AOH1996, MRTX1133 or their combination showing significant loss of clonogenic capacity in the combination-treated cells. **(C)** Western blot of PDAC cells treated with AOH1996, MRTX1133 or their combination for 48 hours, showing pAKT, AKT, pERK and ERK levels in the cellular lysates. **(D)** Bar graph depicting the percentage of cells in each phase of the cell cycle as measured by a PI DNA content analysis assay using flow cytometry. A representative flow plot of each condition is shown on the right. **(E)** Synergy analysis plots using Synergy Finder software showing significant synergy between AOH1996 and AMG 510 (sotorasib) on the left, or with MRTX849 (adagrasib) on the right. **(F)** Western blot of MIA PaCa-2 cells treated with AOH1996, sotorasib or their combination for 48 hours showing pERK and ERK staining.

A subset of pancreatic cancer patients have tumors driven by KRAS^G12C^ mutations and would potentially derive benefit from the FDA-approved KRAS^G12C^ inhibitors AMG-510 (sotorasib) and MRTX849 (adagrasib)^20-22^. Therefore, we tested AOH1996 in combination with sotorasib and adagrasib, which showed strong synergy in G12C mutant cell line MIA PaCa-2 **(Fig. 3E)**. Western blot showed sustained pERK inhibition in MIA PaCa-2 cells treated with sotorasib and AOH1996 **(Fig. 3F)**.

A KRAS-driven human tumor xenograft model was established using the AsPC-1 KRAS^G12D^ xenograft model. Mice received either AOH1996 (100mg/kg, twice a day, 5 days on and 2 days off, orally), or MRTX1133 (15 mg/kg, once a day, 5 days on and 2 days off, orally), or a combination of the two regimens **(Fig. 4A)**. The three treatment regimens were well tolerated, as indicated by body weight measurements **(Fig. 4B)**. The combination treatment of AOH1996 with MRTX1133 was significantly effective at restricting tumor growth compared to single-drug treatments **(Fig. 4C)**. A picture of the excised tumors is shown in **Fig. 4D**. The excised tumors in the combination groups weighed significantly less than the other groups, indicating strong induction of tumor regression by the combination of AOH1996 and MRTX1133 **(Fig. 4E)**. IHC and Western blot were performed to confirm sustained inhibition of KRAS downstream signaling *in vivo* **(Fig. 4F, G)**. Western blot reveals downregulation of pAKT, pERK and Myc indicating strong inhibition of KRAS signaling pathway and reduced proliferation **(Fig. 4F)**. Histology of the excised xenografts stained with hematoxylin and eosin (H&E), Ki67, or pERK antibody, revealed a trend of downregulation of pERK by IHC (**Fig. 4G, S2A**).

**Figure 4.**
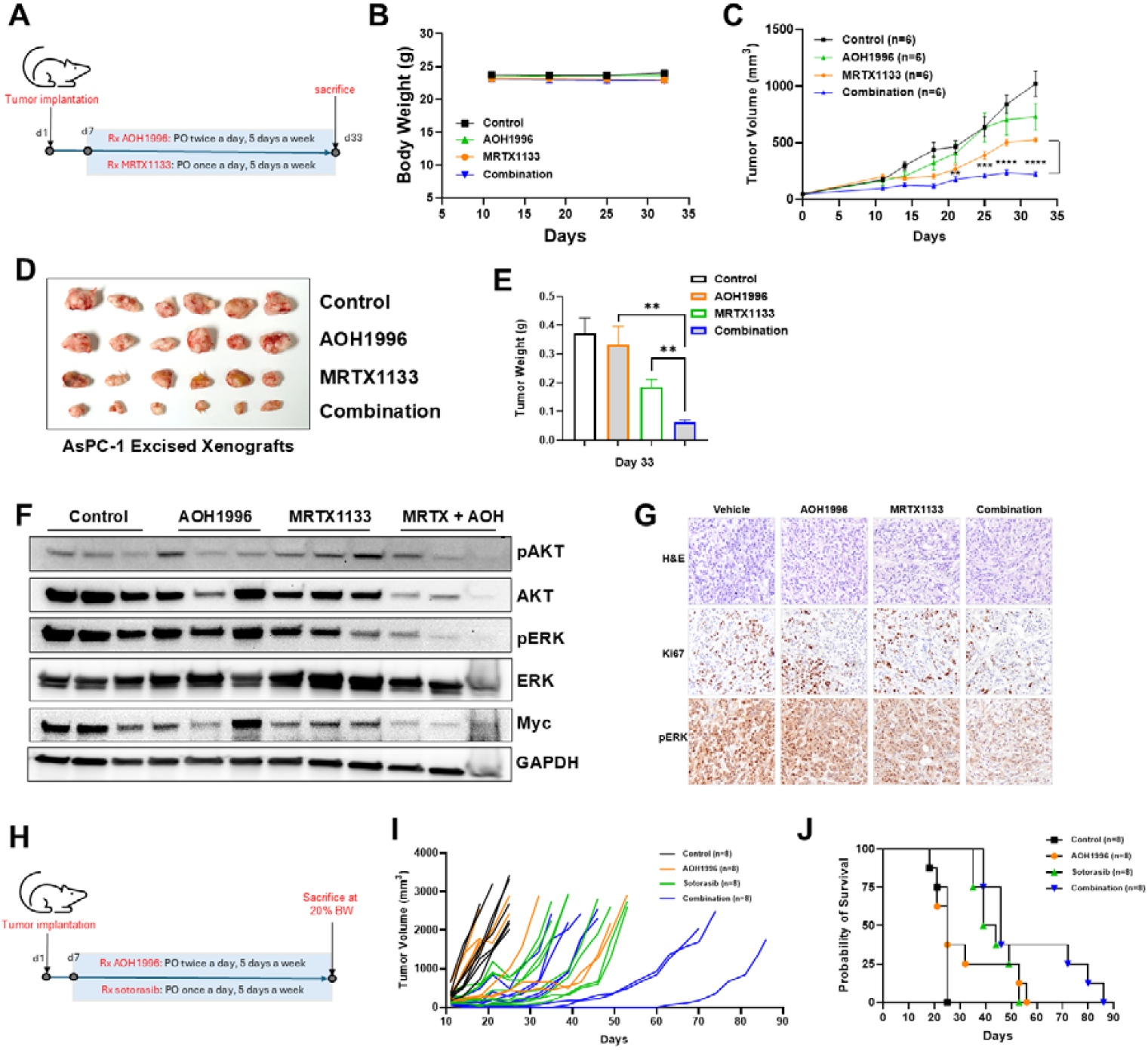
AOH1996 in combination with KRAS^G12D^ inhibitor MRTX1133 shows synergistic anti-tumor activity *in vivo*. **(A)** AsPC-1 xenografts were established subcutaneously in ICR-SCID mice, and were treated with either vehicle, AOH1996, MRTX1133 or a combination of the two drugs. Tumor volumewere followed throughout the treatment for 4 weeks. **(B**) Graph of mice body weight measurement showing no significant loss in body weight throughout treatment (<1%). **(C)** Graph shows tumor volume at corresponding days post-subcutaneous tumor transplantation. **(D)** A picture of the excised tumors i shown. **(E)** Bar graph depicting the weights of the excised mouse xenograft tumors. **(F)** Western blot from three tumor lysates per condition showing pAKT, AKT, pERK and ERK protein levels. **(G)** Images from H&E, Ki67, and pERK IHC FFPE sections of the xenografts. **(H)** ICR-SCID mice were subcutaneously inoculated with MIA PaCa-2 tumor fragments and treated with AOH1996, sotorasib or a combination of the two regimens with 8 mice in each group. **(I)** Spaghetti plot depicting tumor growth in the mice. **(J)** Kaplan-Meier survival plot of the mice described in (H).

In another experiment, mice bearing MIA PaCa-2 subcutaneous xenografts were treated with AOH1996, sotorasib, or a combination of both treatments **(Fig. 4H**). Treatments were well tolerated as shown by body weight measurements **(Fig. S2B)**. Three mice in the combination arm showed a remarkably longer sustained response compared to mice in the other groups **(Fig. 4I)**. Overall, the combination group overall showed a trend towards improved survival **(Fig. 4J)**. Ki67 IHC staining showed a trending downregulation of Ki67 staining **(Fig. S2C)**. Collectively, our *in vivo* studies confirm the synergy we observed *in vitro* between AOH1996 and KRAS-targeted therapies MRTX1133 and sotorasib.

### AOH1996 synergizes with RAS(ON) inhibitor RMC-6236 in cell models, including a primary patient-derived model

In addition to small molecule inhibitors of KRAS, promising KRAS therapeutics include the RAS-selective, tricomplex RAS(ON) inhibitors such as RMC-6236, which is currently in clinical trials for solid tumors with RAS mutations^7^. **Fig. 5A** shows strong synergy between AOH1996 and RMC-6236 in HPAF-II and MIA PaCa-2 cell lines. PDAC patient-derived tumoroid cultures were grown in 3D conditions for combination testing. Primary human tumoroids were isolated directly from the tumor of a patient undergoing surgical resection. The isolated cells are a complex mix that includes tumor and stromal cells, mimicking the patient’s actual tumor and its interaction with the microenvironment. This makes tumoroid models extremely valuable for *in vitro* compound testing for clinical translation. The tumoroids were treated with AOH1996, RMC-6236, or a combination of the two drugs **(Fig. 5B)**. Cell viability assay reveals a significant restriction of tumoroid culture growth in the combination compared to single-agent treatment **(Fig. 5C)**. We also found that the combination results in increased apoptotic cell death in AsPC1 cells. A time-course analysis at 24, 48 and 72 hours revealed an increase in apoptosis at 48 and 72 hours in cells treated with AOH1996 or RMC-6236 and was significantly enhanced in the combination **(Fig. 5D, Fig. S3A, B)**. We also confirmed sustained pathway inhibition in the combination treatment. Combination of RMC6236 and AOH1996 in KRAS^G12D^ mutant cells, AsPC1, showed a sustained inhibition of pERK and pAKT downstream of KRAS **(Fig. 5E)**.

**Figure 5.**
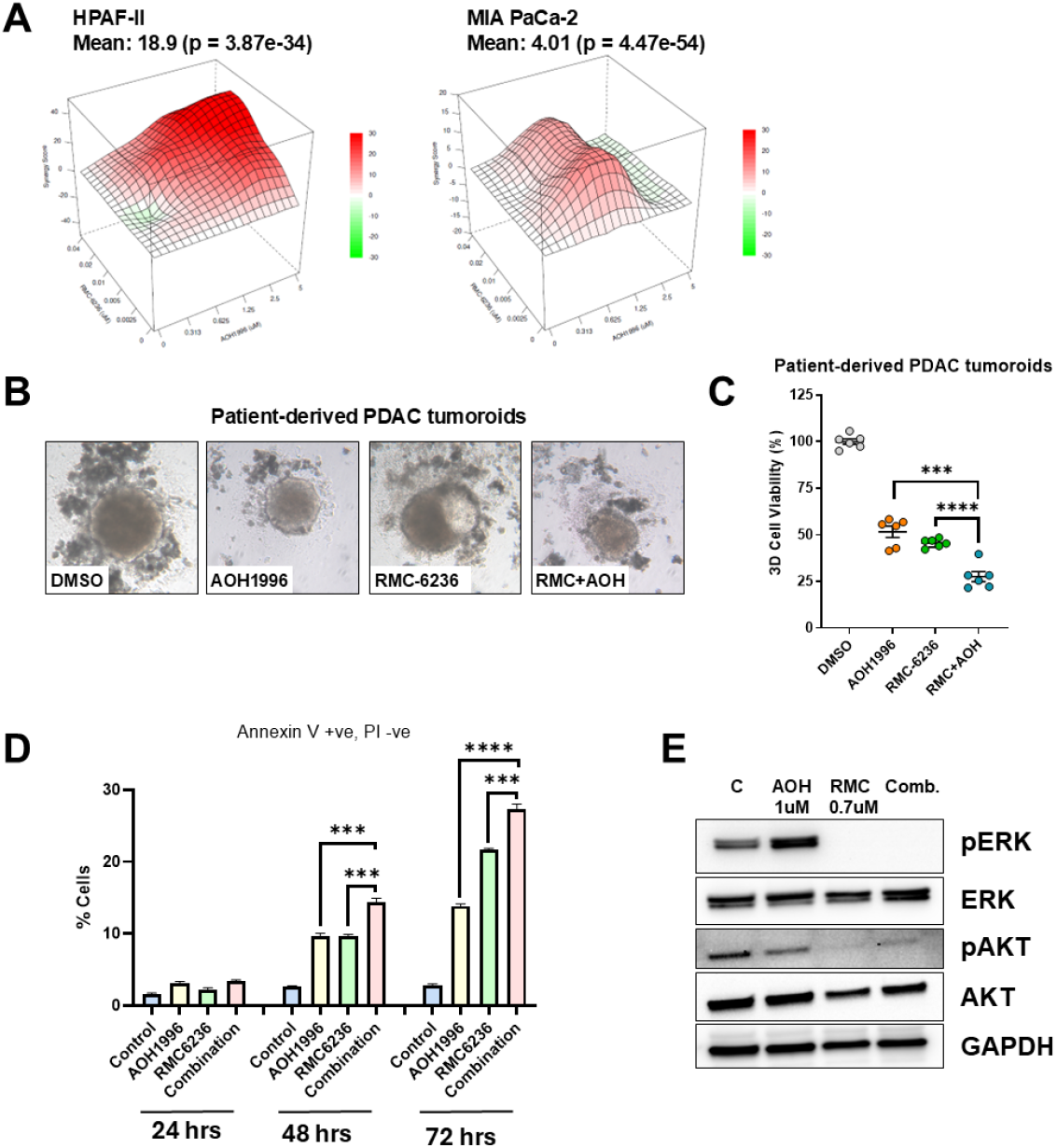
AOH1996 in combination with the RAS(ON) inhibitor RMC-6236 demonstrates synergistic effects in vitro. **(A)** Synergy analysis using Synergy Finder software shows significant synergy between AOH1996 and RMC-6236 in HPAF-II (KRAS^G12D^) and MIA PaCa-2 (KRAS^G12C^) cells. **(B)**Photomicrographs of patient-derived tumoroids containing a complex mixture of primary patient cells grown in 3D conditions and treated with the indicated treatments conditions. **(C)** Plot of the quantification of cellular viability in the tumoroids after the treatments using 3D cell-titer glo. **(D)** Bar graph depicting early apoptotic cells as measured by flow cytometry in cells treated with AOH1996, RMC-6236 or their combination and stained with FITC-Annexin V and PI. Cells that stain for Annexin V only are considered actively apoptotic. **(E)** Western blot of AsPC-1 cell lysates treated with AOH1996, RMC-6236, or their combination for 24 hours, showing the levels of pERK, ERK, pAKT and AKT in different conditions.

## Discussion

RAS is an important oncogenic driver in PDAC and many other malignancies^20^. Exon 2 missense mutations affecting codon 12 are the most prevalent in KRAS mutant PDAC, resulting in a predominantly GTP-bound KRAS and incessant downstream signaling in a ligand-independent manner^23^. Recent medicinal chemistry advances resulted in the FDA approval of inhibitors that target KRAS^G12C^ for lung and colorectal cancer^22,24-26^. Although it has not been approved in PDAC, studies have shown that these inhibitors are safe and effective against PDAC with KRAS^G12C^ mutations^21^. Following the discovery of the switch-II binding pocket within KRAS, inhibitors targeting other mutations, including G12D have been discovered^6^. Moreover, RAS(ON) inhibitors that target the GTP-bound state of both mutant and wild-type KRAS have recently been described^7^. These compounds have been shown to have potent anti-tumor activity in PDAC patients in early clinical studies^27^. Despite these advances, the durability of response to KRAS-targeted inhibitors is limited, and resistance is common^10^. Therefore, there is a critical need for the development of rational combinations to improve the durability of response, thereby reducing resistance and improving the clinical utility of these agents.

PCNA acts as a hub protein for DNA replication and repair pathways^15^. This makes it an essential protein for cancer cell proliferation and survival, as it allows tumor cells to tolerate increased DNA damage and cellular stresses. Previous studies have shown that AOH1996 inhibits PCNA by a unique trapping mechanism by stabilizing the interaction of PCNA with the largest subunit of RNA polymerase II, RPB1, causing its degradation and preventing the resolution of transcription replication conflicts (TRC)^14^. A recent study has shown that this mechanism is important for the activity of AOH1996 in pancreatic ductal adenocarcinoma^28^. Treatment of PDAC cells with AOH1996 stalled the progression of replication forks and induced TRC-related DNA damage^28^. Initial clinical evidence also shows that AOH1996 is nontoxic and results in tumor shrinkage in patients with PDAC^28^.

In this study, we show that PCNA inhibition using AOH1996 synergizes with novel KRAS inhibitors in both G12D and G12C mutant models. Transcriptome studies on cells treated with AOH1996 demonstrated that the treatment has an impact on MAPK signaling pathway. We examined KRAS pathway inhibition in cell lines as well as xenograft tumors and found a sustained inhibition of pERK and pAKT as a result of the combination treatment. In a patient-derived primary tumoroid model, we found that the combination of AOH1996 with RMC6236 produced a strong anti-tumor response. Our *in vivo* testing showed arrested tumor growth in the combination treatment. In our KRAS^G12C^ model, we observed a heterogeneous response to the treatment, with three exceptional responders to the combination treatment. ICR-SCID mice, which were used as a model in this experiment, have an outbred background. Therefore, individual mice demonstrate genetic diversity that increases experimental variability, thus making outbred ICR-SCID mice a more relevant model to human subjects compared to an inbred model. The outbred ICR strain exhibits high genetic diversity, similar to human populations, while the SCID mutation allows for xenograft implantation, making this model extremely valuable for studying drug sensitivity and response. Our findings in this experiment underscore the importance of identifying biomarkers of response to allow for optimal clinical translation of this combination.

Overall, this study demonstrates that targeting PCNA with AOH1996 is a promising strategy for PDAC, with potent single-agent activity and robust synergy with KRAS inhibitors across in vitro, ex vivo and in vivo models. The combination of AOH1996 with a KRAS-targeted drug is a valuable strategy for KRAS-mutant PDAC, warranting further evaluation and translational development of the combination regimens for patients with PDAC.

## Acknowledgements

The Biobanking and Correlative Sciences Core and Molecular Imaging and Core at Karmanos Cancer Institute are supported, in part, by NIH Center Grant P30 CA022453 to the Karmanos Cancer Institute at Wayne State University. Asfar Azmi’s lab acknowledges the support from SKY Foundation Inc, U CAN-CER VIVE Foundation and Perry Family Foundation. SFB acknowledges support from the DeRoy Testamentary Foundation. The funders had no role in the design of the study, in the collection, analyses, or interpretation of data, in the writing of the manuscript, or in the decision to publish the results.

