## Supplemental Materials for "PCNA Inhibition Enhances the Antitumor Activity of KRAS-Targeted Therapies in Pancreatic Cancer"

Supplementary figures and table

Table S1

| PRIMER TARGET | PRIMER SEQUENCE |
|---|---|
| MAP2K6 | F- 5'-AAACGGCTACTGATGGATTTGG-3' |
| | R- 5'-CAGTGCGCCATAAAAGGTGAC-3' |
| CMPK2 | F- 5'-GTACCTCCTTTATTCCTGAAGCC-3' |
| | R- 5'-ATGGCAACAACCTGGAACTTT-3' |
| SPDEF | F- 5'-CACCTGGGGCGATTCACTAC-3' |
| | R- 5'-GATGAGTCCACCTCGCTGTC-3' |
| BARX2 | F- 5'-CAGCGAGTCAGAGACGGAAC-3' |
| | R- 5'-CTGAGCCAAGTCCAACCTGT-3' |
| NUPR1 | F- 5'-TCGGAGGTGGAGGCCG-3' |
| | R- 5'-TCACCAGTTTCCTCTCGTGC-3' |
| BEST1 | F- 5'-CTGGGCTTCTACGTGACGC-3' |
| | R- 5'-TTGCTCGTCCTTGCCTTCG-3' |
| EFR3B | F- 5'-CGTCCTATCACCGGAGCTATG-3' |
| | R- 5'-GCCTTTGATGCCTGACATTCG-3' |

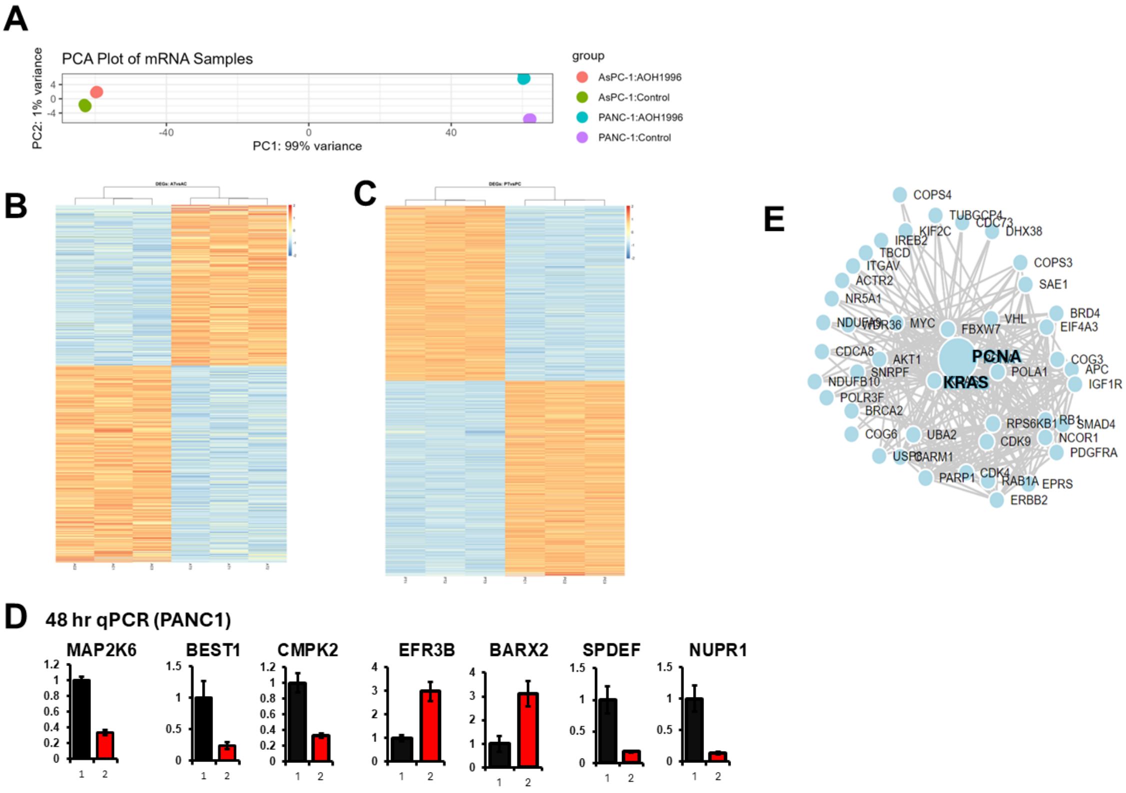


**Supplementary Figure 1. Related to Figure 2**

**(A)** PCA plot of all samples used in RNA-seq analysis. **(B)** Heatmap of differentially expressed genes (DEGs) in AsPC1 cells treated with AOH1996. **(C)** Same as (B) in PANC1 cells. **(D)** Validation of some genes from RNA-seq data using RT-qPCR. **(E)** Synthetic lethality analysis for PCNA using Slorth database (University of Sussex Bioinformatics Lab).

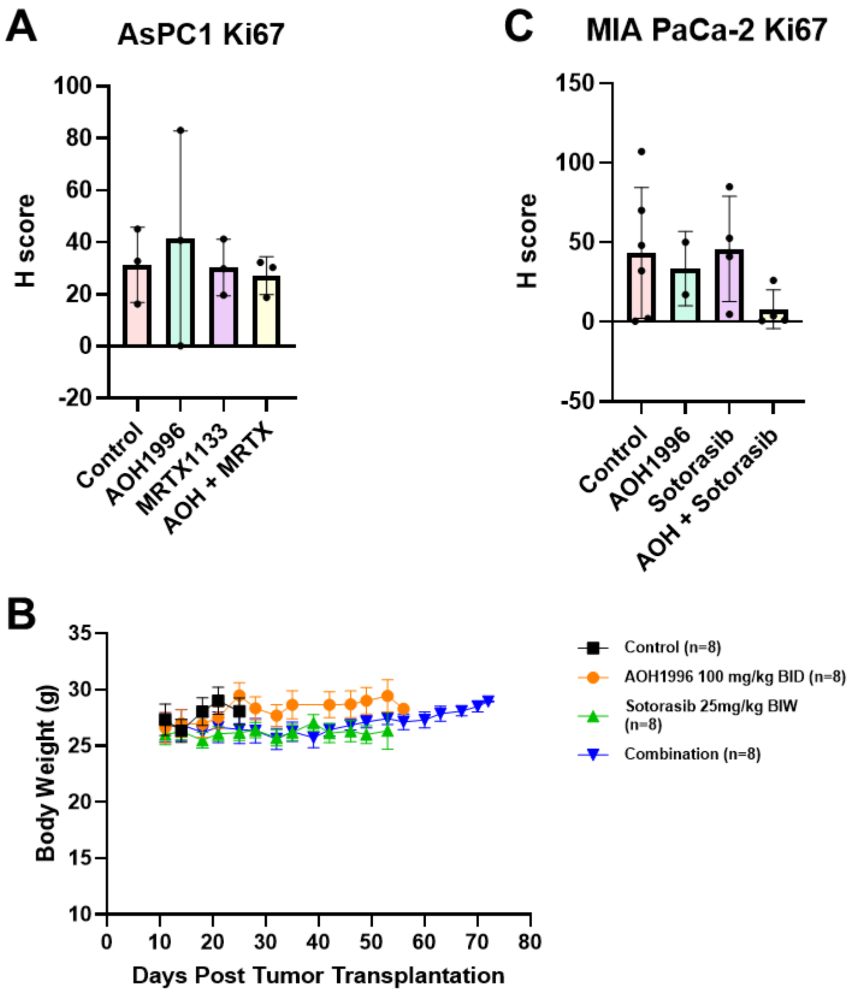


**Supplementary Figure 2. Related to Figure 4**
**(A)** H score quantification of Ki67 stain in AsPC1 xenograft FFPE sections. **(B)** ) Graph of body weight measurements of mice described in Fig. 4 H-J. **(C)** Ki67 H scores in MIA PaCa-2 xenografts.

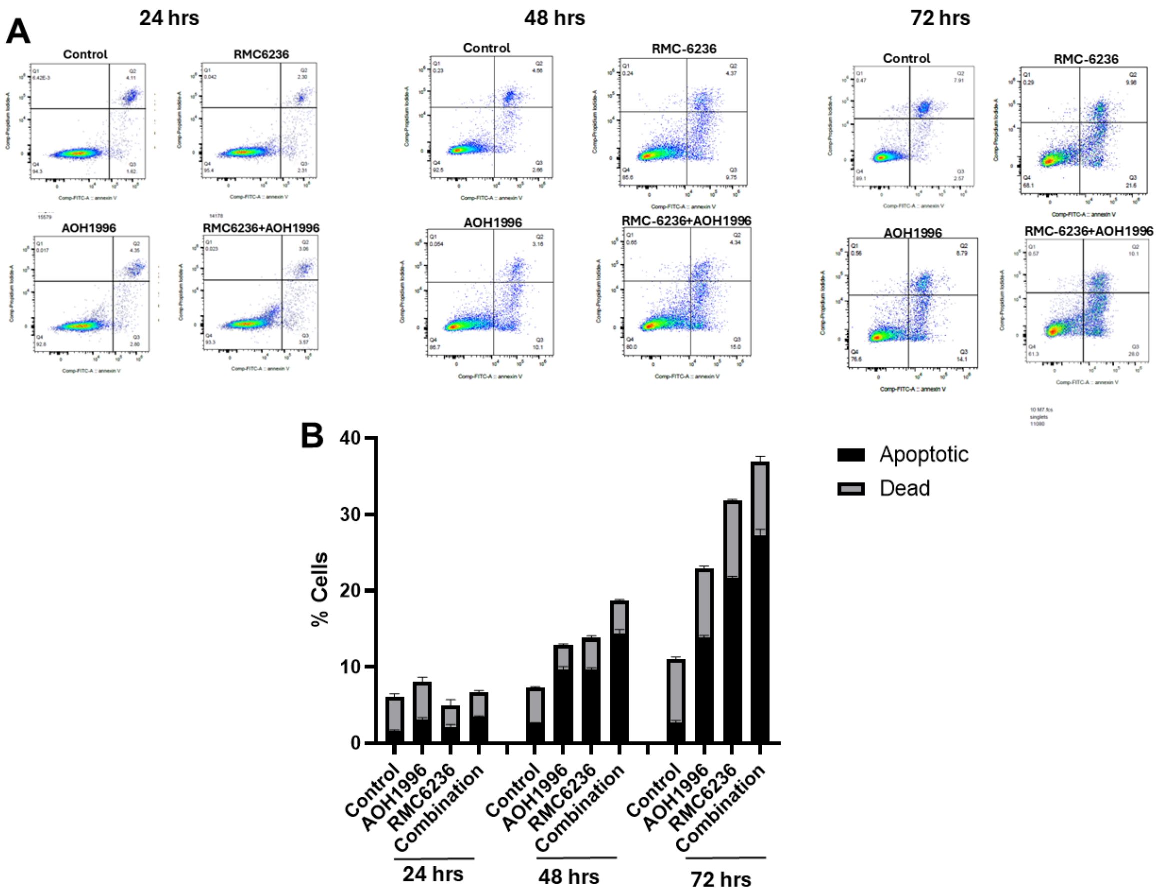


**Supplementary Figure 3. Related to Figure 5**

**(A)** Representative flow cytometry scatter plots of AsPC-1 cells treated with vehicle control, RMC-6236, AOH1996 or their combination, and stained with FITC-Annexin V and PI stains. **(B)** Bar graph depicting early apoptotic and late apoptotic (dead) cells as measured by flow cytometry as shown in (A). Data represents 3 replicates.
